# Reinforcement learning for adaptive control of phenotypically heterogeneous bacterial populations

**DOI:** 10.1101/2025.11.21.689767

**Authors:** Josiah Kratz, Zihang Wen, Shiladitya Banerjee, Oana Carja

**Author notes:** Equal corresponding authors (ordered alphabetically). E-mail(s). Contributing authors. Equal contribution (ordered alphabetically) (Josiah Kratz, Zihang Wen).

## Abstract

Bacterial populations display extraordinary resilience to antibiotic stress, driven by diverse physiological states that allow some cells to persist and later repopulate. This phenotypic heterogeneity, amplified by environmental fluctuations, undermines the effectiveness of conventional fixed-dose treatment regimens. To address this challenge, we introduce a reinforcement learning (RL) framework that discovers adaptive treatment strategies using only experimentally accessible, population-level measurements. The RL agent learns to infer the hidden physiological state of the population and leverages this knowledge to maintain control even under conditions not encountered during training. Moreover, when granted control over nutrient availability, an important driver of physiological change often overlooked in antibiotic treatment protocols, the agent consistently drives population extinction, surpassing adaptive protocols based solely on drug dynamics. This computational framework offers a powerful, data-driven approach for designing adaptive treatment strategies to counter the growing threat of antimicrobial resistance.

## Introduction

Although structurally simple, bacteria display remarkable phenotypic adaptability. By reconfiguring their sensory networks and regulatory pathways, they can dynamically reprogram their physiology to survive in hostile and unpredictable environments [1, 2]. Even within genetically identical populations, substantial variation in gene expression, resource allocation, and stress responses frequently emerges [3–6]. These phenotypic states can further persist as a form of epigenetic “memory” of prior stress, leading to faster adaptation to future challenges [7–11].

This remarkable plasticity also makes bacterial populations notoriously difficult to control [12–14]. As in other rapidly evolving biological systems such as cancer cell populations [15], standard fixed-dose treatments often fail, selecting for phenotypically resistant subpopulations that evade drug action and enable population regrowth [16–18]. In response, recent years have seen growing interest in adaptive treatment strategies: dynamic therapeutic protocols that continuously adapt treatment decisions in anticipation of evolving population dynamics [19–22]. However, most existing approaches rely on detailed mechanistic knowledge of system dynamics or require information (like real-time subpopulation composition) that is difficult to obtain in practice [19, 23, 24]. Furthermore, many of these models overlook non-genetic heterogeneity, the wide range of physiological states that demand feedback-guided control under environmental uncertainty.

Reinforcement learning (RL) provides one such model-free framework capable of operating robustly under uncertainty. An RL agent iteratively refines its strategy through evaluative feedback to maximize long-term outcomes, even when knowledge of the underlying biological system is incomplete. Recent studies have shown that RL can identify adaptive dosing policies despite uncertainty, and these approaches have already been applied to therapeutic decision-making in several contexts, including bacterial and cancer cell populations [22, 25–28].

However, existing RL implementations rely on simplified, low-dimensional models that neglect stochasticity in physiological adaptation and focus exclusively on drug administration, overlooking other key ecological variables [22, 27, 28]. Nutrient availability, a major determinant of bacterial physiology, is a particularly important omission, as it strongly influences cellular fitness and drug susceptibility [29–33]. Little is also known about how well these algorithms generalize beyond the specific environmental conditions on which they are trained.

Here, we address these limitations by introducing an RL framework that incorporates bacterial phenotypic stochasticity and jointly controls both antibiotic exposure and nutrient availability. Specifically, we employ a deep Q-learning (DQN) algorithm [34, 35] to manipulate a virtual bacterial population whose intracellular dynamics capture both single-cell behavior and population-level responses across diverse, time-varying environments [33]. By reshaping bacterial physiology through targeted adjustments to nutrient levels and drug dosing, and relying solely on population growth rate measurements, the agent learns a policy that accelerates short-term population growth to render cells more susceptible and increase the proportion of sensitive cells before applying treatment. This strategy ultimately reduces population growth and enhances long-term drug efficacy. Notably, consistent population extinction emerges only under this dual-control paradigm, an outcome not achieved by prior drug-only approaches.

A key innovation is the coupling of RL with a mechanistic, multiscale cell model that connects gene expression dynamics to population-level outcomes. This integration not only enables optimal control of a complex system using experimentally accessible measurements, but also provides mechanistic insight into why specific strategies succeed. Such interpretability is essential for moving adaptive control strategies toward clinical application. Moreover, we show that our framework remains effective despite unobserved environmental perturbations, successfully infers hidden nutrient fluctuations, and generalizes to conditions not encountered during training. Taken together, our findings highlight nutrient modulation as a crucial lever for adaptive therapy and establish RL-based physiological control as a powerful approach for suppressing rapidly evolving cell populations.

## Results

### Challenges of controlling evolving, phenotypically heterogeneous bacterial populations

To illustrate the challenges of controlling microbial populations through environmental modulation, we begin by characterizing the behavior of evolving bacterial populations using a multiscale computational model that simulates individual cells as they grow, divide, and die over time [33]. The model explicitly links intracellular dynamics to the external environment, with nutrient availability and antibiotic concentration as the primary environmental variables governing population growth and survival.

Each cell in the population is defined by a set of intracellular state variables that describe its physiological condition. These include the mass fractions of major proteome components (such as ribosomal, metabolic, and stress-response proteins) and the concentration of antibiotic-induced damage accumulated over time. When this damage surpasses a critical threshold, the cell is assumed to undergo death (Fig. 1A). The change in a single cell’s internal state is described by a system of stochastic differential equations:

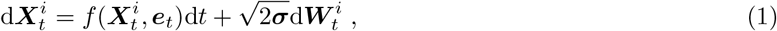

where 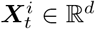 is a vector representing the intracellular state of cell *i* at time *t*, and *f* describes how the cell responds to its environment and evolves in time (see Methods for the full description of the multiscale model). The vector ***e***_*t*_ = [*b*_*t*_, *c*_*t*_] denotes the external environmental parameters (antibiotic concentration and nutrient availability), ***σ*** represents diffusion coefficients capturing intrinsic noise, and 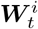 is a Wiener process in ℝ^*d*^.

**Fig. 1.**
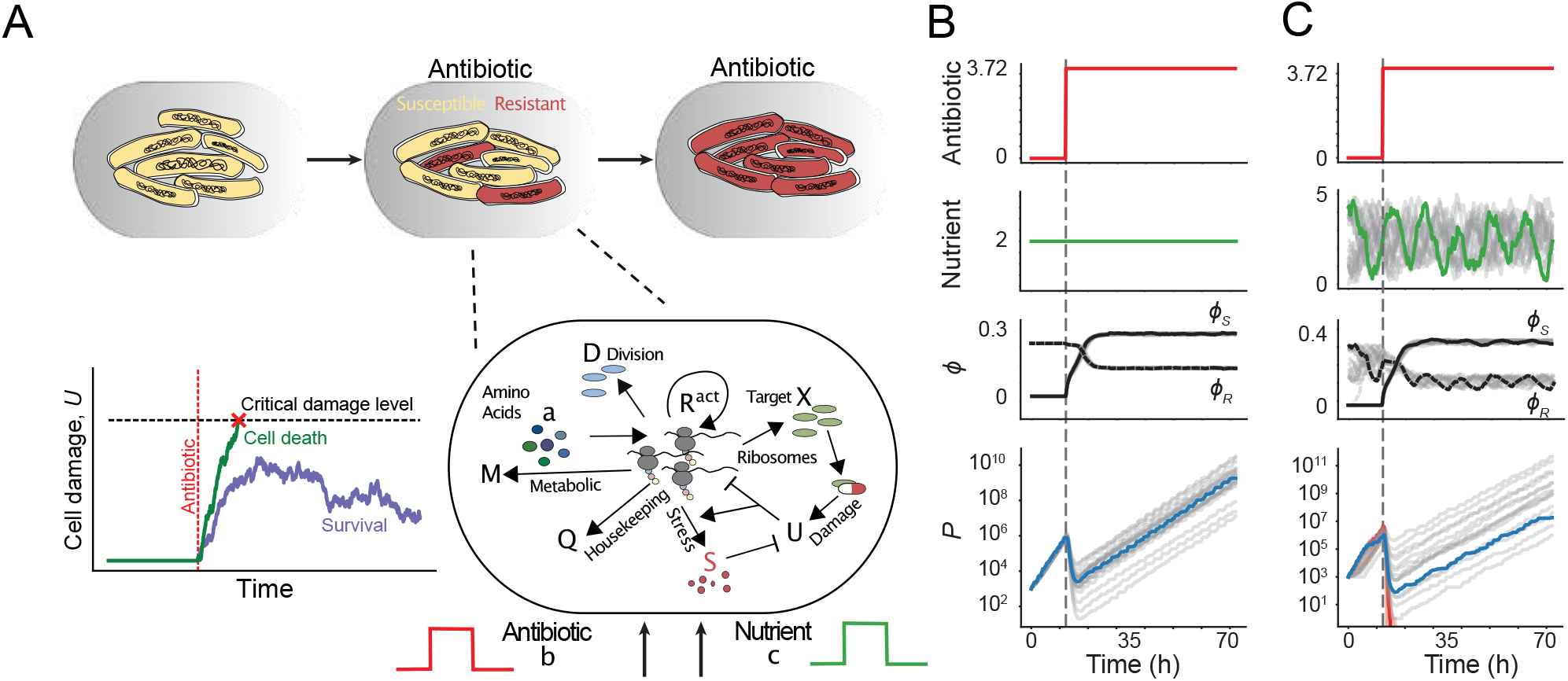
**A** Schematic depicting resistance emerging within a bacterial population. In response to antibiotic application, some cells stochastically accumulate damage to a critical level and die, while others develop resistance by altering their physiological state. This process is modeled by simulating stochastic changes to gene expression and damage accumulation in response to environmental changes, for each individual cell. **B** Under constant nutrient conditions, all populations eventually develop resistance and survive when antibiotics are applied (gray line), with stochastic population dynamics. Resistance development is mainly driven by increased expression of proteins involved in combating stress response (*ϕ*_*S*_). **C** In stochastic pulsatile nutrient conditions, populations show even greater heterogeneity, punctuated by divergent survival outcomes: some populations develop resistance (representative trajectory shown in blue), while others go extinct (red trajectory). See Methods for simulation details.

By explicitly linking the extracellular environment to intracellular dynamics, our model enables simulation of both single-cell and population-level behavior across a broad range of time-varying conditions. Under constant nutrient conditions, antibiotic exposure triggers cells to reallocate proteomic resources toward the production of stress-response proteins that repair or remove damage (Fig. 1B). This adaptive response decreases the susceptibility of surviving cells to subsequent doses, reducing the death rate and increasing the minimum inhibitory concentration of the population [33], allowing cells to recover from initial inhibition and eventually resume growth. The system’s behavior changes dramatically when nutrient availability fluctuates over time: some populations successfully adapt and persist, while others fail to recover and ultimately go extinct (Fig. 1C).

This stochastic model highlights a central challenge in microbial population control, namely, that no single fixed treatment strategy is consistently effective. Even with a detailed mechanistic description of population dynamics, the inherent variability in outcomes across microbial populations makes the control problem fundamentally complex and resistant to simple, predetermined intervention protocols.

### RL-guided optimization of antibiotic strategies in constant nutrient environments

Given that constant antibiotic application yields unreliable outcomes, we sought to develop a dynamic, feedback-guided control strategy that relies only on experimentally accessible measurements, such as population growth rate. To ensure broad applicability in real-world scenarios, where accurate mechanistic models of the biological system are often unavailable, this approach must also be model-free. To address these challenges, we employ a model-free deep Q-learning (DQN) framework [35] (Fig. 2A). Unlike model-based optimization approaches, the DQN agent learns an optimal control policy directly through iterative interaction with the system and and evaluation of outcomes, without requiring prior knowledge of its internal dynamics. This property makes DQN particularly well-suited for stochastic, partially observed, and multiscale biological systems. Handling partial observability is particularly important because, in most experimental and clinical settings, only population-level measurements like growth rate are available. Although these measurements offer valuable insight into system behavior, they do not uniquely reveal underlying physiological states, such as proteome allocation or intracellular damage. As a result, the agent must infer effective control strategies from limited and aggregated information.

**Fig. 2.**
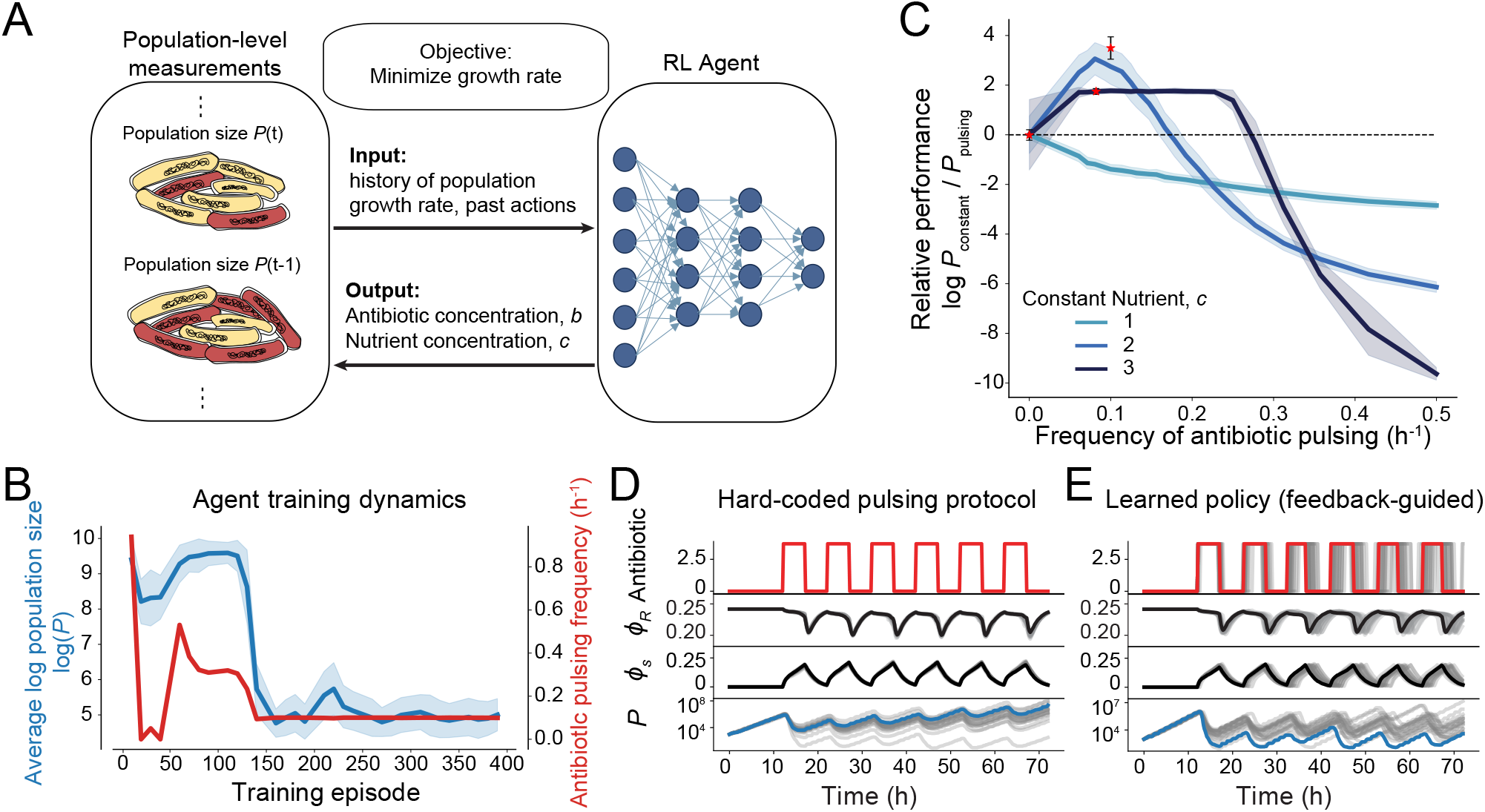
**A** Schematic of approach. An RL agent is trained to minimize the discounted cumulative cost, given by the log population growth rate, log(*P*_*t*_*/P*_*t*−*τ*_ )*/τ* . To learn the optimal strategy, the agent iteratively chooses actions (defined by nutrient and antibiotic concentrations) and receives feedback following model simulation to generate the next population-level measurements. **B** Agent training dynamics. The agent training protocol consisted of 400 episodes, with performance evaluations conducted every 10 episodes using 100 test runs. Performance was quantified by the average value of the logarithm of the population size (blue line, shading denotes one standard deviation). Simultaneously, agent behavior was tracked, as given by the average antibiotic pulsing frequency (red line). After approx. 175 episodes, both performance and learned pulsing frequency stabilized, indicating convergence. **C** Feedback-guided learned policies outperform hard-coded pulsing. Agents were trained in different constant nutrient environments and then evaluated and compared to a range of hard-coded pulsing protocols with different frequencies, where performance is measured by the log ratio of the final population size under constant drug application (*P*_constant_) and under drug pulsing (*P*_pulsing_). Thus, policies with relative performance above the horizontal dotted line outperform constant drug application. Learned policies (red stars) matched or outperformed hard-coded ones (colored lines). Shading denotes one standard deviation from the average of 100 simulations. The dashed line denotes constant antibiotic application. **D, E** Representative population and physiological state dynamics for cells subjected to a hard-coded antibiotic protocol with fixed pulsing frequency (D) compared to the RL agent’s feedback-guided policy (E), for 100 realizations. See Methods for agent hyperparameter details.

In the DQN framework, our objective is to learn a policy *π* that maps the system *states* **x**_*t*_ to control *actions* **a**_*t*_, that modify the environment to steer the system toward a desired physiological state. Formally, this mapping is expressed as, **a**_*t*_ = *π*(**x**_*t*_). Here 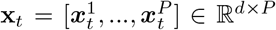 represent the state variables of all cells in a population of size *P*, which can change dynamically as cells grow, divide, or die.

The *optimal* policy, *π*^∗^, is defined as the policy that minimizes an objective function quantifying treatment performance. In our formulation, this objective corresponds to driving the population toward extinction by minimizing the expected cumulative discounted cost, represented by the population growth rate:

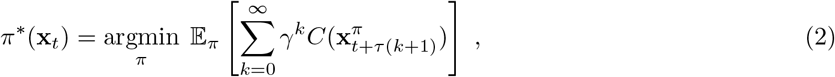

where *C* is the instantaneous cost and *γ* ∈ [0, 1) is the discount factor, which determines how myopic the agent is (see Methods for more details). To more closely mimic real-world dosing protocols, we discretize the control problem: the system evolves continuously over a fixed interval *τ*, during which the chosen action is held constant. The cost function *C* is based on experimentally measurable quantities, specifically, the logarithmic change in population size over time, and assigns a large reward when the population goes extinct. Finally, to improve learning performance in a continuous state space under partial observability, the DQN agent’s input is augmented with a history of past actions and observed growth rates, providing temporal context that helps the agent infer hidden system dynamics (see Methods, Supplementary Figure S1).

We first evaluated our reinforcement learning approach by training agents to learn a ‘bang-bang’ control policy [36, 37], where the agent chooses whether to apply or withhold antibiotic treatment at each decision point (*a*_*t*_ ∈ {0, *b*_max_}). This binary decision structure simplifies the control problem while mimicking realistic clinical scenarios. Agents were trained across a range of constant nutrient environments, and, in all cases, they learned to accept short-term costs by withholding treatment temporarily, in order to preserve long-term drug efficacy. Throughout the training process, agents demonstrated robust rapid learning and stabilized their pulsing frequency patterns to successfully control the growth of the bacterial population (Fig. 2B).

To assess the optimality of the learned policies, we compared the RL agent’s performance against a set of hard-coded periodic antibiotic pulsing protocols, across a range of periods (Fig. 2C). Because nutrient availability strongly alters protein production and population death rates, which together ultimately influence bacterial susceptibility, the optimal pulsing frequency varied across conditions. In every case, the RL-derived policy matched or exceeded the performance of the hard-coded pulsatile protocols (Fig. 2C), demonstrating its ability to discover near-optimal solutions without prior knowledge of system dynamics.

Comparison with the best hard-coded protocols further highlights the advantages of feedback-guided control. Pulsing maintains antibiotic efficacy by allowing cells to revert to a more susceptible physiological state, as measured by low values of *ϕ*_*S*_, during drug-free intervals (Fig. 2D). However, unlike static strategies, the RL agent adapts its pulsing frequency dynamically in response to real-time growth-rate feedback, which reflects changes in population susceptibility. This enables context-specific control policies that more effectively suppress bacterial growth (Fig. 2E).

These results show that our RL framework can exploit bacterial stress-response dynamics to design adaptive, optimized treatment protocols. By tailoring dosing in real time based on system feedback, the RL approach provides a powerful alternative to traditional predetermined strategies.

### Agent-controlled nutrient availability increases drug efficacy and can drive bacterial populations extinct

Having shown that our agent can optimize antibiotic timing to outperform hard-coded dosing strategies, we next explored whether it could dynamically leverage nutrient availability to reshape bacterial susceptibility and enhance drug efficacy [14].

To establish a baseline, we first evaluated two hard-coded dual-control protocols that jointly regulate antibiotics and nutrient conditions: *Feast* and *Famine* (Fig. 3A, B). Motivated by evidence that nutrient-rich environments increase bacterial susceptibility to antibiotics [14], the Feast protocol pairs a nutrient upshift with antibiotic treatment. Although this approach initially reduces population size, it consistently fails to achieve population extinction (Fig. 3A). Conversely, the Famine protocol combines antibiotic treatment with a nutrient downshift to induce starvation stress. This strategy performs better overall, achieving extinction in a subset of cases, though outcomes vary from complete eradication to continued proliferation (Fig. 3B).

**Fig. 3.**
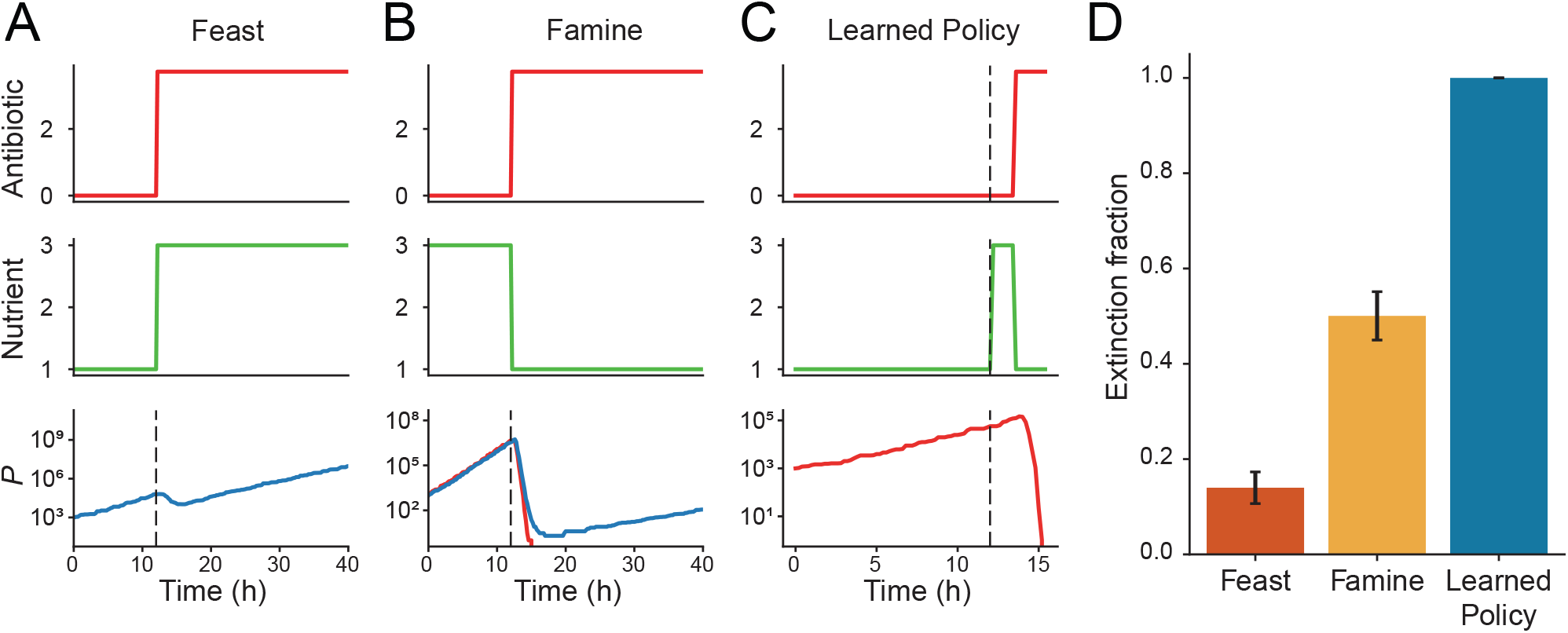
Learned policy improves antibiotic efficacy through context-dependent nutrient modulation. **A** Example trajectory of the ‘Feast’ protocol, in which antibiotics are applied (top, red) concurrently with rich nutrients (middle, green), regardless of initial condition. This approach results in an initial phase of population death followed by recovery and proliferation (bottom, blue). **B** Example trajectory of the ‘Famine’ protocol, in which antibiotics are applied (top, red) concurrently with a nutrient-poor environment (middle, green), regardless of initial condition. This approach results in heterogeneous population survival outcomes, with some realizations resulting in extinction (bottom, red), while others result in survival (bottom, blue). **C** Example trajectory of the RL agent’s learned policy, in which antibiotics and nutrients are dynamically adjusted based on the population’s initial physiological state, consistently resulting in population extinction. **D** Quantification of the success of each approach, as given by the extinction fraction after 72 hours of simulation, across 100 realizations. Unlike the hard-coded policies, the agent is able to achieve extinction in every case. (Details about the simulation can be found in Methods, Section 2.)

To identify more effective strategies, we expanded the agent’s action space to include nutrient modulation: **a**_*t*_ = [*b*_*t*_, *c*_*t*_], where *b* and *c* again represent antibiotic concentration and nutrient availability, respectively. The agent learned an adaptive policy that resembled the Famine strategy but incorporated several key refinements (Fig. 3C). When populations were initialized in nutrient-poor environments, the agent deliberately provided nutrients while withholding antibiotics, allowing growth to shift cells into a more susceptible physiological state. Once susceptibility increased, the agent simultaneously applied antibiotics and restricted nutrients, inducing starvation stress. This coordinated, dual-stress approach maximized bacterial vulnerability.

Critically, the feedback-guided nature of the RL approach provided a decisive advantage over fixed protocols. When initial treatment attempts failed to achieve extinction, the agent adapted by iteratively reapplying the dual-control sequence (Supplementary Fig. S2), ultimately eliminating the population in all trials (Fig. 3D). In contrast, fixed protocols either failed consistently (Feast) or succeeded only sporadically (Famine) (see Supplementary Fig. S2). These results demonstrate that feedback-driven dual control significantly improves treatment reliability and efficacy compared with predetermined strategies.

### Inference of hidden nutrient dynamics enables adaptive control in dynamic environments

By manipulating the nutrient environment, the RL agent was able to enhance drug efficacy beyond what could be achieved by simply increasing the dosage. However, in many real-world scenarios, environmental factors like nutrient availability cannot be directly controlled. For example, bacteria colonizing a host encounter dynamic nutrient conditions driven by physiological processes, such as periodic changes in blood glucose following meals [38, 39]. This variability necessitates control strategies robust to unobserved environmental fluctuations, requiring agents to infer both physiological state and underlying nutrient dynamics solely from growth rate observations.

Motivated by this clinically relevant context, we introduced unobserved stochastic fluctuations in the nutrient environment with a predefined period *T* . Specifically, we modeled nutrient dynamics using a Langevin equation with cyclic drift *u*_*t*_ and mean-reversion timescale *τ* :

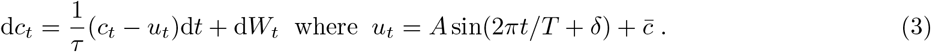

Here, the constants 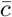 and *A* denote the predefined average nutrient value and amplitude, respectively. The RL agent was trained on bacterial populations exposed to these fluctuating conditions with random initial phase offsets δ, and without direct access to nutrient values. Thus, it was required to infer nutrient dynamics and their impact on susceptibility solely from growth observations and action history.

Despite these constraints, the trained agent successfully drove populations to extinction (Fig. 4A). Our analysis revealed a sophisticated decision-making process: following antibiotic application, bacteria upregulated stress proteins as a defense mechanism (Fig. 4A, B). Simultaneously, nutrient availability declined, increasing resistance. In response, the agent learned to strategically pause antibiotic treatment, waiting for stress protein levels (*ϕ*_*S*_) to decrease and nutrient availability to recover, thereby restoring susceptibility (Fig. 4A, B). The agent then resumed antibiotic application, repeating this cycle until population eradication (Fig. 4A). This demonstrates the agent’s ability to integrate bacterial physiological state, environmental context, and treatment history to optimize therapeutic interventions in uncertain, dynamically changing environments.

**Fig. 4.**
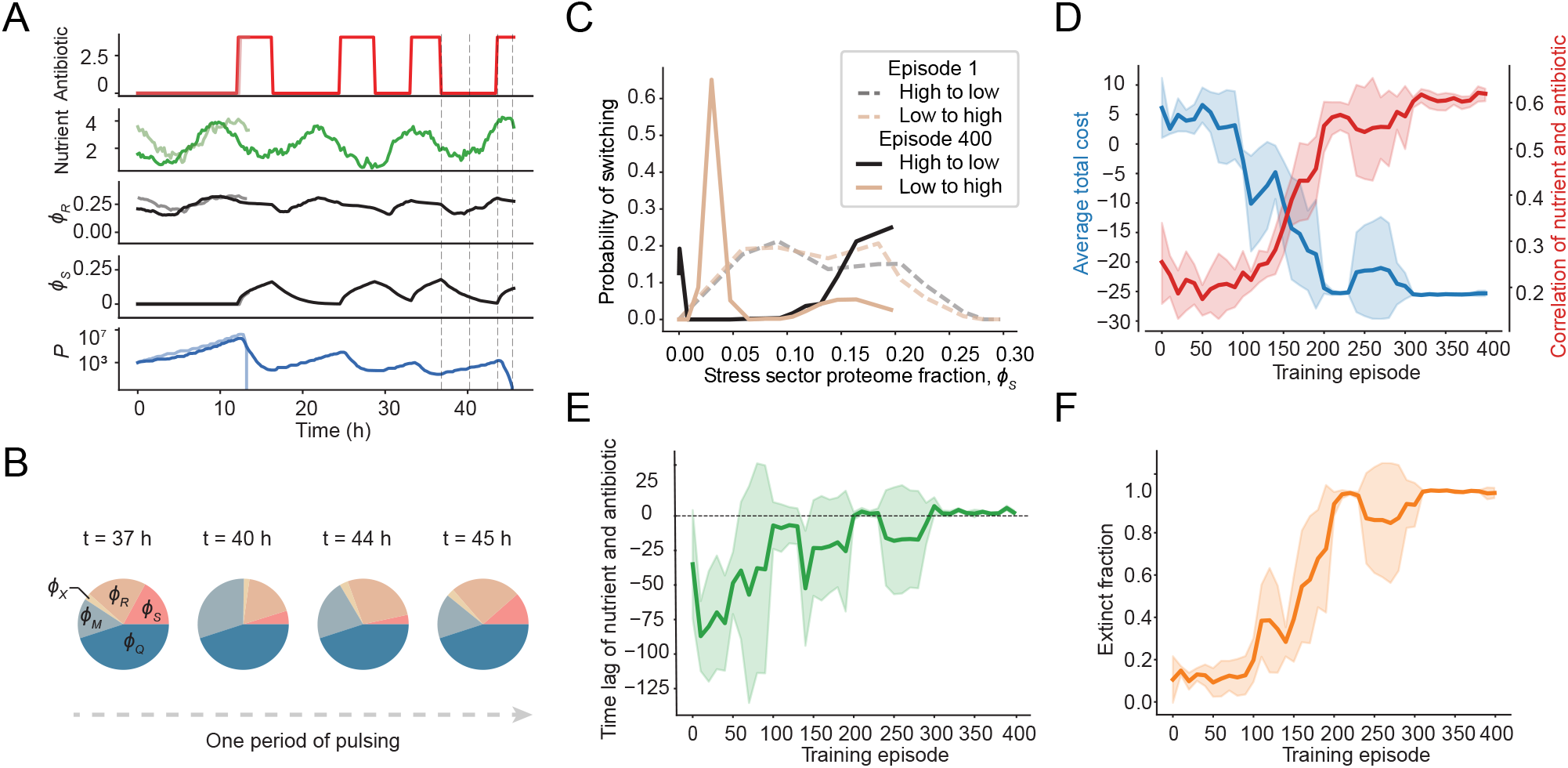
Agent’s learned protocol successfully infers bacterial physiological state despite unobserved nutrient fluctuations. **A** Two representative evaluations of an agent trained in a scenario in which the nutrient environment is fluctuating cyclically, but the nutrient level is unobserved. Plots of the learned policy (top, red), unobserved stochastic nutrient environment (second from top, green), average ribosomal and stress protein expression (third and fourth from top, respectively, shown in black and gray), and the resulting population trajectories, show that, for populations which survive the initial antibiotic application, the agent can successfully drive extinction through a pulsing protocol in which the pulse timing is determined by the resulting physiological state. **B** Population-average proteome allocation pie charts corresponding to different time points in the plot in **A** (denoted by the blue dashed lines) showing that pulsing stops when cells are more resistant (caused by high values of *ϕ*_*S*_), and resumes when cells are in a more susceptible state (low value of *ϕ*_*S*_). **C** Probability of switching between antibiotic application (high) and absence (low) and vice versa as function of average stress protein expression across training. Probability of switching was quantified by binning switching events into equally-sized bins and quantifying the fraction. At episode 1 (no training), the probabilities of switching from low to high and vice versa are almost identical, and depend little on the stress sector proteome fraction (dashed lines), which is a measure of cell susceptibility. After training, at episode 400, the agent learns to apply the antibiotic (low to high, solid gold line) with high probability when stress protein expression is low, and to withhold antibiotic (high to low, solid black line), when stress protein expression is high. **D** Average training dynamics. Average total cost (blue line), as quantified as the average log population ratio across an episode, and average cross-correlation between nutrient and antibiotic timeseries (red line) across 400 training episodes. Correlation between nutrient and antibiotic increases with performance, indicating the agent is able to infer the underlying nutrient state through training. **E** Throughout the course of training, the time lag of the maximum cross-correlation between nutrient and antibiotic timeseries stabilizes at a small positive value, indicating that the agent learns to respond to nutrient fluctuations when determining when to apply antibiotic. **F** Average extinction fraction throughout training. Extinction fraction stabilizes near one at the same time that lag and cross-correlation stabilize, indicating that learned policy performance is dependent on successfully inferring the environmental state.

To further characterize the agent’s behavior, we compared the switching probabilities between high and low antibiotic concentrations before and after training (Fig. 4C). Prior to training, the policy behaved essentially randomly, with antibiotic application decisions showing no relationship to the underlying physiological state of the population. Post-training, the agent consistently applied antibiotics when stress-protein levels were low (indicating increased susceptibility) and withheld treatment when they were high. Moreover, the agent’s decisions became strongly correlated with background nutrient availability, showing successful inference of nutrient fluctuation patterns (Fig. 4D).

A positive temporal lag between nutrient availability and antibiotic application confirms the agent’s actions were informed by previously inferred environmental states rather than immediate conditions (Fig. 4E). This behavior remained robust across different nutrient oscillation periods (Supplementary Fig. S3), highlighting the agent’s capacity to integrate past environmental information with current observations to guide treatment decisions. Training metrics confirmed the effectiveness of this adaptive strategy: both average treatment cost and extinction fraction improved significantly after training (Fig. 4D, F). Moreover, the feedback-driven policy substantially outperformed hard-coded pulsing protocols (Supplementary Fig. S4).

Together, these results show that RL agents can operate effectively in realistic, partially observed, and dynamic environments by developing sophisticated inference capabilities that integrate multiple indirect information sources.

### Learned policies generalize to unseen environments

The RL agent’s learned policy must generalize effectively to environmental dynamics not encountered during training. To assess this capability, we trained several single-environment ‘specialist’ agents (on either constant or oscillatory nutrient profiles of varying periods) and a multi-environment ‘generalist’ agent (trained on a mixture of all profiles) (Fig. 5A). We hypothesized that the generalist would outperform the specialists in unseen conditions, as broader training distributions are typically associated with improved generalization.

**Fig. 5.**
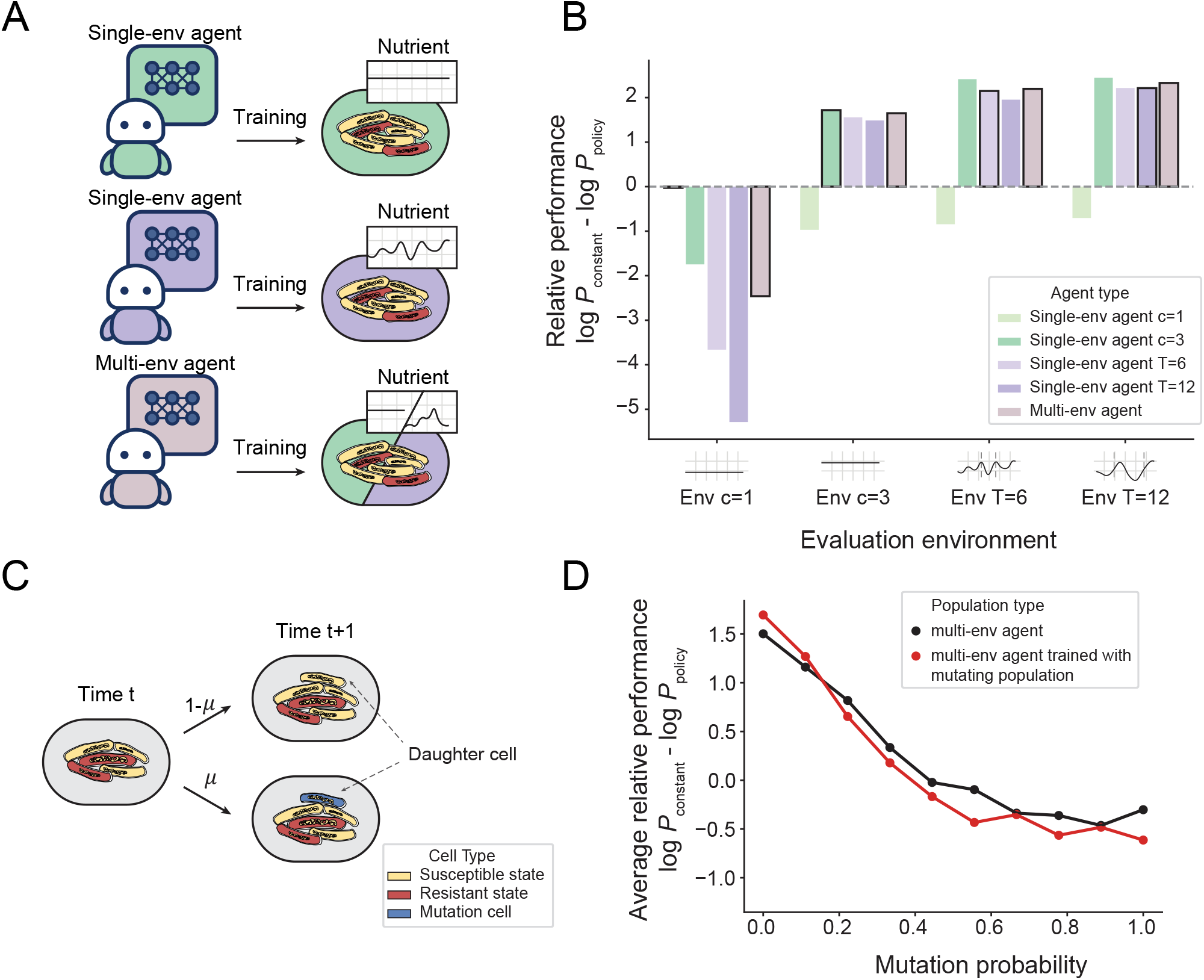
A Schematic depicting RL agent training protocol. Single-env ‘specialist’ agents were trained in a single environment type: either constant nutrient conditions (nutrient availability *c* ∈ {1, 3}) or stochastic oscillatory conditions (time period *T* ∈ {6, 12}). The Multi-env ‘generalist’ agent was trained across multiple environments (nutrient availability *c* ∈ {1, 2, 3, 4}, time period *T* ∈ {6, 12, 18, 24}), with the nutrient condition randomly chosen at the start of each episode. **B** Relative performance, given by the log ratio of the final population size under constant drug application (*P*_constant_) and under the learned policy (*P*_policy_), for 100 trials in four different test environments. **C** Schematic of the mutation scenario. At each cell division, a daughter cell mutates with probability *µ*, acquiring a new value of 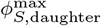 following Eq. (4). **D** Average log population reduction compared to constant drug application across different experiments for 100 trials as a function of mutation probability *µ*. Black dots denote the performance of the Multi-env agent detailed in **5A** and **5B**, which was trained for 400 episodes in different nutrient environments in the absence of mutations. Red dots denote the performance of the Multi-env agent, which was trained for 500 episodes with a bacterial mutation probability of *µ* = 0.3.

Surprisingly however, the specialist agents performed as well as, and in some cases better than, the generalist when evaluated on previously unseen environments (Fig. 5B). This result suggests that specialists learned robust control strategies by internalizing a representation of cell susceptibility that was largely invariant to specific nutrient dynamics. To further validate this robustness, we evaluated agent performance in additional stochastic environments generated by Gaussian processes without periodic structure. The agents again maintained strong performance across a range of correlation lengths and amplitudes (Supplementary Fig. S5), demonstrating effective generalization beyond the conditions encountered during training.

### Learned policies remain effective against mutating populations, but generalization degrades under extreme non-stationarity

The robust performance demonstrated against diverse environmental dynamics suggests that the learned policies capture context-transcending principles of physiological susceptibility. However, a fundamental challenge for long-term control is resilience to genetic evolution. While physiological state dictates drug susceptibility on short timescales [14, 40], mutations are the primary mechanism of antibiotic resistance over longer timescales [41]. To test generalization under these conditions, we extended our multi-scale model to incorporate mutations that alter resistance.

Mutations can influence susceptibility by changing drug–target binding, or indirectly, by modulating global gene expression through regulatory networks [42, 43]. Here, we focused on the latter, modeling mutations as an independent multiplicative process in which individual cell parameters change once per division (Fig. 5C). Specifically, the maximum allocation to stress response proteins for cell 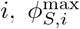, mutates with probability *µ* ∈ [0, 1] according to

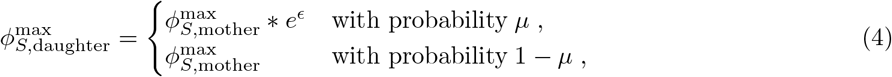

where *E* ∼ *𝒩* (0, *σ*^2^). These mutation-driven parameter changes further complicate the control problem by distorting the relationship between observed growth rate and underlying physiological state.

To test the limits of generalization in this setting, we tested performance across a range of mutation rates, many of which are much higher than those seen in actual bacterial populations [44]. As shown in Fig. 5D, agent performance declines significantly as mutation probability increases, demonstrating that generalization of policies trained on non-mutating populations degrades with the introduction of mutations. At extremely high mutation rates (*µ >* 0.4), performance plateaued near zero, meaning agents performed comparably to constant drug application. However, for populations with biologically-relevant mutation probabilities (*µ «* 0.1), agent performance was well above the constant application plateau, thus substantially outperforming the naive constant-application protocol over the duration tested (3 days). These findings suggest that RL-driven control remains advantageous in biologically relevant regimes over intermediate timescales (days to weeks), but performance will degrade as mutations accumulate within the population.

Surprisingly, training an agent directly on simulations that included mutations did not lead to better control strategies; performance matched that of agents trained on non-mutating populations (Fig. 5D). This outcome is due to mutations introducing non-stationarity into the system, violating a key assumption of the Bell-man equation that underpins our Q-learning algorithm (see Methods and Supplementary Note 1). While our approach is robust to stochastic nutrient dynamics, rapidly mutating populations with non-stationary transition dynamics present a fundamental limitation of current RL-based methods. Addressing this limitation, for example, by modifying the algorithm to incorporate contextual information, represents an important direction for future work [45].

## Discussion

We developed a model-free deep Q-learning framework that learns adaptive antibiotic treatment strategies using only experimentally-accessible population growth measurements, without requiring knowledge of underlying cellular states. By leveraging these limited observations, our reinforcement learning approach successfully infers the heterogeneous physiological states of bacterial populations to achieve effective control. This directly addresses a fundamental challenge in antimicrobial therapy: the diverse adaptive behaviors and stochastic proteome reallocation that enable bacterial populations to recover even under sustained treatment [33, 46–48]. The feedback-guided policies learned through RL consistently outperform conventional treatment protocols by dynamically adjusting interventions in response to real-time population and environmental conditions.

The trained agents discovered control strategies that substantially exceeded the performance of conventional approaches across diverse scenarios. In constant nutrient environments, agents learned to temporarily withdraw antibiotics, accepting short-term population growth, to restore drug susceptibility, achieving better control than optimized hard-coded pulsing protocols. When given control over both antibiotics and nutrients, the agent developed a dual-control strategy: initially providing nutrient-rich conditions to induce susceptibility, followed by coordinated application of nutritional and antibiotic stress to drive population extinction. This approach succeeded where fixed protocols either failed completely or worked only stochastically, demonstrating how these methods can identify non-obvious protocols that exploit complex physiological relationships.

Crucially, the agent maintained effectiveness even with unobserved environmental fluctuations. When trained under stochastic nutrient oscillations, the agent successfully inferred both physiological states and underlying nutrient dynamics solely from growth observations and treatment history. The learned policy strategically applied antibiotics when inferred stress protein levels were low and withheld treatment when conditions favored resistance. Furthermore, agents trained in a single environment generalized robustly to unseen nutrient dynamics, performing comparably to those trained across multiple conditions. This strongly suggests that the learned policies capture fundamental, context-transcending relationships between observable dynamics and physiological susceptibility.

Our approach demonstrates the unique benefit of testing RL learned policies on mechanistic biological models. This coupling enables mechanistic insight into why specific, seemingly “black box” strategies succeed or fail, which is essential for clinical translation and offers new opportunities for biological discovery when integrated with experimental validation. While our agents exhibited robust generalization across a wide range of environmental conditions, introducing mutation-driven resistance evolution revealed important limitations. When mutations that alter stress protein expression capacity were incorporated, modeling global gene expression changes that affect drug susceptibility, agent performance decreased with increasing mutation probability. This occurs because mutations introduce non-stationary dynamics that violate the core Markov assumption underlying our Q-learning framework, distorting the relationship between observed growth rates and underlying physiological states throughout treatment.

Addressing this challenge represents a key opportunity for future algorithmic work, incorporating techniques from the nonstationary RL literature [49–55], such as meta-learning for rapid adaptation [53], and context-aware Markov decision processes for inferring evolutionary states [51]. Beyond algorithmic development, experimental validation of learned strategies and establishing regulatory frameworks for adaptive treatment algorithms remain essential for clinical translation. As antimicrobial resistance continues to evolve through both physiological adaptation and genetic mutation, computational approaches must similarly evolve to maintain effective treatment options.

## Methods

### Virtual bacterial population model

In this study, we utilize a previously developed and validated model for bacterial growth and death dynamics in dynamic nutrient and antibiotic environments from Ref. [33]. We briefly summarize the dynamics here.

The state of cell *i* at time *t* is described by its intracellular composition 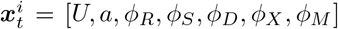 (Fig. 1A). We do not simulate the dynamics of housekeeping proteins (*Q*), as the mass fraction is assumed to remain constant. To grow, a cell imports and converts nutrients to amino acids, with mass fraction *a* (not to be confused with the RL agent’s action ***a***_*t*_ in the main text), via metabolic proteins, with protein mass fraction *ϕ*_*M*_ . Amino acids are consumed by translating ribosomes, with mass fraction *ϕ*_*R*_, to synthesize all proteins, including themselves. As a result, *ϕ*_*R*_ sets the cellular growth rate, specifically 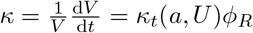, where *κ*_*t*_(*a, U* ) quantifies the translational efficiency, which is a function of *a* and *U* (see Ref. [33] for more details). Here, *ϕ*_*D*_, *ϕ*_*X*_, and *ϕ*_*S*_ represent the mass fractions of division, target, and stress proteins, respectively, which collectively determine population growth and division rates (see below for more details). The dynamics of each sector are given by:

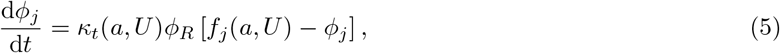

where *j* = [*R, S, D, X, M* ] and *f*_*j*_(*a, U* ) denotes the fraction of total cellular protein synthesis flux devoted to sector *j* and can be a function of *a* and/or *U* (see Ref. [33] for a complete description of the functions *f*_*j*_). The dynamics of the amino acid mass fraction are given by the difference in the metabolic and translational fluxes, specifically

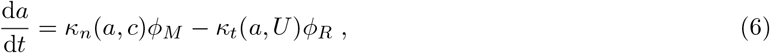

where *κ*_*n*_(*a, c*) quantifies the rate of nutrient import, and is a function of *a* and nutrient quality *c*.

Cell division is triggered when a cell produces a sufficient number of *D* proteins, i.e. *D*(*t*_division_) = *D*_0_, at which point it divides into two daughter cells which inherit the same intracellular composition as the mother. The dynamics of *D* are given by:

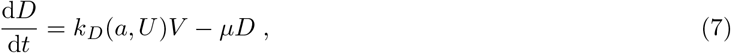

where *V* is the cell volume, *µ* is the degradation rate of protein *D*, and *k*_*D*_(*a, U* ) is the division protein synthesis rate per unit volume (see Refs. [56] and [33] for more details).

Critically, each cell maintains a damage concentration *U* caused by drug-target binding. Cell death occurs when damage accumulation exceeds a critical level *U*_0_, such that *U* (*t*_death_) = *U*_0_. The stochastic dynamics of *U* are given by:

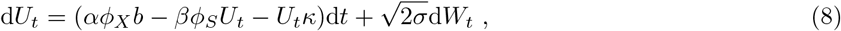

where *b* is the antibiotic concentration, which produces damage at a concentration specific rate *α* when bound to its target protein *X*. Damage is actively removed by stress proteins *S*, at a rate *β*, and is also diluted via growth with growth rate *κ*.

Finally, we can generically describe this process as:

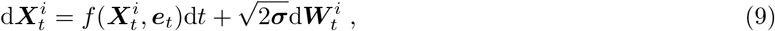

where ***e***_*t*_ = [*b*_*t*_, *c*_*t*_]. We present this form of the dynamics in the main text.

### Reinforcement learning for control

The goal of stochastic control can be formulated as learning a feedback-guided protocol *π* (referred to as a *policy* in RL) mapping the coordinates in phase space describing the system **X**_*t*_ (*states*) to values of the control inputs **a**_*t*_ ∈ **e**_*t*_ (*actions*), which alter the environment to achieve some desired physiological state of the system. Thus, **a**_*t*_ = *π*(**X**_*t*_). Here we write **X**_*t*_ ∈ ℝ^*d×P*^ to denote the state variables of all cells in a population of size *P* (where *P* can change with time as cells die and/or divide). Thus we can rewrite Eq. (9) as:

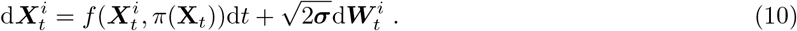

The *optimal* policy maximizes (or minimizes) an objective function representing its overall performance across a given time horizon.

We hope to learn control policies that reduce population growth and drive extinction over long timescales. In this setting with no clear time horizon, we can define an infinite horizon control problem where the expected cumulative discounted cost starting from population state **x**_*t*_ (referred to as the *value function*) is given by:

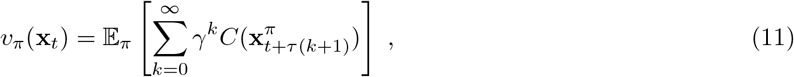

where *C* is the instantaneous cost and *γ* ∈ [0, 1) is the discount factor. Discounting is needed mathematically for the sum to converge in the infinite horizon limit (assuming *C* is bounded), but also allows for tuning of how myopic the control policy is. In practice, we treat *γ* as a hyperparameter that is tuned to achieve the best performance. Critically, here we assume that **x**_*t*_ evolves continuously for a constant duration *τ* over which the action is fixed after being determined by the initial value **x**_*t*_ (i.e. **a**(*s*) = *π*(**x**_*t*_) for *s* ∈ [*t, t* + *τ* ]). Then **x**_*t*+*τ*_ is obtained by solving Eq. (10) with the initial condition **x**_*t*_. This discretization better mimics real-world control applications, but also introduces lag into the feedback protocol. However, in practice as long as *τ* is small, this lag does not pose serious challenges to performance.

There have been many different techniques developed to find the policy *π* which maximizes Eq. (11) (cf. [57] and [58] among others for an overview).

Here, we employ a specific class of model-free deep reinforcement learning algorithms based on Q-learning. Q-learning lifts the expected return *v* so that it depends on a given state-action pair (**x**_*t*_, **a**_*t*_) and policy *π*. We can define the *action-value* function for policy *π*, or Q-function, as:

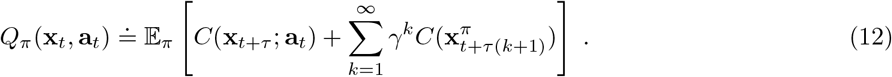

The expectation is taken over trajectories initialized at **x**_*t*_, subjected to the action **a**_*t*_ for a duration *τ*, and thereafter following policy *π*. In other words, the Q-function describes the total discounted reward of taking action **a**_*t*_ in state **x**_*t*_, assuming all future actions are given by policy *π*. The optimal Q-function, *Q*^∗^, is then the one in which all future actions are given by the optimal policy, *π*^∗^. If *Q*^∗^ is known, then computing the optimal policy is straightforward, as it is simply given by:

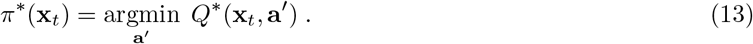

However, when encountering a new system we seek to control, we must first learn *Q*^∗^. If the transition dynamics and reward function are known, and the state space Ω is small, *Q*^∗^ could in principle be computed via dynamic programming. However, in practice when dealing with real biological systems, we rarely have access to good dynamical models of the system, and must solely learn *Q*^∗^ through interaction and data collection from the biological system itself. *Q*^∗^ can be approximated by noting that [58, 59]:

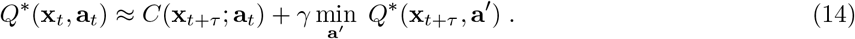

Beginning with some initial guess, we can iteratively learn an approximation for the optimal Q-function, 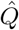, by sampling actions and recording state transitions, then using the update rule:

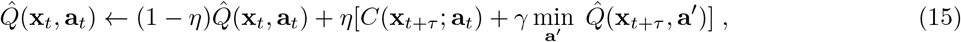

where *η* ∈ [0, 1] is the learning rate.

When dealing with continuous state spaces, as is the case when working with state variables that describe biological systems, the state space is too large to learn an action-value for each state. However, deep neural networks can be used to approximate the Q-function and interpolate action-values without needing to visit every state [60]. In this approach, we parametrize the Q-function with network parameters *θ*. Learning then occurs through an iterative procedure in which actions are initialized randomly, but transition over time to greedy action selection. After an action is executed, an experience tuple (**x**_*t*_, **a**_*t*_, *C*_*t*_, **x**_*t*+*τ*_ ) is stored in a replay buffer ℬ and used to make parameter updates to the network. Specifically, experience tuples are sampled randomly from the replay buffer to compute the Bellman residual, the squared difference between the right and left-hand sides of the optimal Q-function (Eq. (14)):

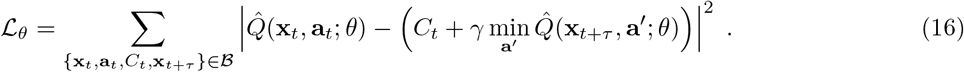

The network parameters *θ* are then updated via mini-batch gradient descent with respect to ℒ_*θ*_. Utilizing a neural network enables learning of highly-performant, complex policies even when the state-space is large.

As is common in value-based algorithms, we use an additional target network to reduce the effect of bootstrapping during training. The target network (denoted with parameters 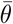) has its parameters frozen during training, and updated periodically via weighted averaging with the online network parameters 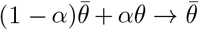. We additionally use the double DQN approach to mitigate the over-estimation bias in standard DQN [61]. We tune over several hyperparameters (whose values we list in the Hyperparameters section) to optimize performance.

Unlike other optimization methods, such as optimal control or model-based RL, the optimal action-value function *Q*^∗^ is learned directly through interaction, and does not require access to any model of the environment, it simply learns through sampling. This is beneficial for several reasons. Firstly, it allows for optimization of stochastic, multiscale models which are hard to directly optimize. Secondly, in an experimental setting, we typically only have access to population-level measurements.

Furthermore, although we seek to control a bacterial system governed by Eq. (9), due to experimental constraints we typically only have access to observations which partially describe the system. For example, in this work we focus on population growth rate. Although growth rate is certainly a quantitative measure of the physiological state of the population, multiple underlying physiological states (proteome allocation and damage concentration values) can result in the same growth rate, thus it does not fully describe the underlying state of the system. In addition, this measure does not capture population heterogeneity. More formally, this can be described by introducing a function *h* : ℝ^*d×P*^ → ℝ which maps the true underlying state to observations, such that *O*_*t*_ = *h*(**x**_*t*_, **a**_*t*_), where here again *P* is the number of cells.

This partial observability means that the same observation can correspond to different underlying states, and thus breaks the Markov assumption used to formulate the Q-learning approach [57]. A straightforward way this challenge can be addressed is by augmenting the observation with some summary *g* of the past history of the controlled trajectory (*H*_*t*_ = (**a**_*t*_, …, **a**_*t*−*kτ*_, *O*_*t*_, …, *O*_*t*−*kτ*_ )), such that 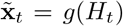. The resulting state 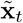 does not have to be the same as the true underlying state of the system **x**_*t*_, but it needs to be able to restore (at least approximately) the Markov property to the dynamics. A complete history of all past actions and states (*g* being the identity function) would be sufficient to recover the Markov property, but this is computationally inefficient, as our description of the state 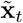 would grow unboundedly with time. One straightforward approach which we employ is to augment the current observation with the past *k* actions and observations. This approach is analogous to delay embedding for systems identification [62].

### A note on the choice of cost function

The expected cumulative discounted cost is given by:

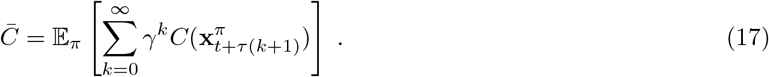

As we seek to minimize the long-term population growth rate of the entire bacterial population and drive extinction, we choose the cost to be instantaneous population growth rate. Assuming population growth is approximately exponential (i.e. *P* (*t*) = *P*_0_*e*^*Ct*^), then this can be estimated by the log ratio of the population size between decisions, divided by the decision interval: *C*(**x**_*t*_) = log(*P*_*t*_*/P*_*t*−*τ*_ )*/τ* for *P*_*t*_ *>* 0. When *P*_*t*_ = 0, the cost function above is undefined, thus we instead give a large numerical reward which incentivizes the agent to learn extinction protocols. The resulting instantaneous cost function is then:

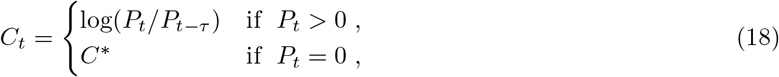

where *C*^∗^ is a large negative value such as 1 − *e*5.

Importantly, the discount factor *γ* alters the control problem, thus altering the optimal solution. If *γ* = 0, then 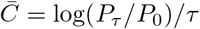, thus the agent is only rewarded for minimizing population growth over the first decision interval without regard for population size at longer times. In contrast, if *γ* = 1, then 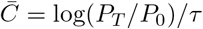 (where *T* denotes episode length). Thus, in the absence of discounting and for large values of *T*, the agent could, in theory, allow the cell population to become extremely large at intermediate times before driving extinction, which is highly suboptimal from a clinical perspective. As a result, we choose *γ* = 0.99 to reward the agent for decreasing population growth over long timescales, while also preventing the agent from allowing the cell population to increase substantially over intermediate timescales.

### Simulation hyperparameters

All simulation episodes were initialized with a 12-hour warm-up period during which cells grew freely without control inputs (in order to remove potential effects from cell initialization), followed by 60 hours during which the agent had control. All mathematical details and parameters for the multi-scale model of bacterial growth can be found in Ref. [33].

### RL model hyperparameters

Each Q-network was parameterized by a four-layer multilayer perceptron (MLP) with 64 neurons each and ReLU activation [63]. During training, we used the Adam optimizer [64] with learning rate 1 *×* 10^−4^. We used a buffer size of 1 *×* 10^6^, batch size 512, and reward discount factor *γ* = 0.99. The network update rate was *α* = 0.005.

The history length used for partial observability in all models was set to 30 time steps. A comparison of performance across different history lengths is shown in Supplementary Fig. S1.

### Bootstrapping for agent-controlled nutrient availability

To quantify uncertainty in the extinction fraction for the agent-controlled nutrient condition, we used a bootstrap procedure. We first generated 100 independent simulations. We then performed 1000 bootstrap resamples, each consisting of 100 draws with replacement from these simulations. Each resample produced one estimate of the extinction fraction, yielding 1000 bootstrap estimates in total. These 1000 values were used to compute the mean and variance of the extinction fraction.

## Code availability

Custom scripts were used for the simulations and data analyses. All code is available on GitHub at https://github.com/quantumyeasthacker/RL-for-bacterial-control. All packages used for analysis and visualization are open-source.

## Acknowledgments

S.B. acknowledges support from the National Institutes of Health (NIH R35 GM143042), and the Shurl and Kay Curci Foundation. O.C. acknowledges support from the National Institutes of Health (NIH R35 GM147445) and the National Science Foundation (NSF CAREER Award 2442397).

## Competing interests

The authors declare no competing interests.

## Notes

https://github.com/quantumyeasthacker/RL-for-bacterial-control

### Competing Interest Statement

The authors have declared no competing interest.

