## Supplemental for "Reinforcement learning for adaptive control of phenotypically heterogeneous bacterial populations"

### 1 Including mutation events make the population dynamics nonstationary

We model the mutation process as a series of discrete events which occur every cell replication cycle. Each mutation event can either increase or decrease the fraction of the proteome devoted to stress protein production, $\phi_S^{\max}$. Let us first consider a single cell which grows and divides. Let $Y_0 = \phi_S^{\max,i}(t=0)$ denote the value of $\phi_S^{\max}$ for cell $i$, before any mutation events. The first mutation event is modeled as:

$$Y_1 = Y_0\epsilon_1 \quad \text{where } \log(\epsilon_1) \sim \mathcal{N}(\mu, \sigma^2) \ . \tag{1}$$

Similarly, the next mutation event leads to:

$$Y_2 = Y_1\epsilon_2 = Y_0\epsilon_1\epsilon_2 \ . \tag{2}$$

Extending this to $t$ sequential mutation events we obtain:

$$Y_t = Y_0 \prod_{i=1}^{t} \epsilon_i \ . \tag{3}$$

Letting $X_t = \log Y_t$, we obtain the sum:

$$X_t = X_0 + \sum_{i=1}^{t} z_i \quad \text{where } z_i \sim \mathcal{N}(\mu, \sigma^2) \ . \tag{4}$$

Calculating the mean and variance of $X_t$ yields:

$$\mathbb{E}(X_t|X_0) = X_0 + \mu t \ , \tag{5}$$

$$\mathrm{Var}(X_t|X_0) = \sigma^2 t \ . \tag{6}$$

Because $X_t|X_0 \sim \mathcal{N}(X_0 + \mu t, \sigma^2 t)$ and $Y_t = e^{X_t}$, we can then write:

$$\mathbb{E}(Y_t|Y_0) = Y_0 \exp(\mu t + \sigma^2 t/2) \ , \tag{7}$$

$$\mathrm{Var}(Y_t|Y_0) = Y_0^2[\exp(\sigma^2 t) - 1]\exp(2\mu t + \sigma^2 t) \ . \tag{8}$$

Clearly the resulting distribution is nonstationary, as both mean and variance change with time.

If one variable in a multivariable stochastic process is nonstationary, then the entire process is considered nonstationary, as the joint statistical structure changes over time. Thus, introducing mutations into our previously stationary intracellular dynamics makes them nonstationary. Importantly, here we have considered the dynamics of a single cell in the absence of antibiotics. When antibiotics are applied, the population is under strong selection, further accelerating the physiological transformation of the surviving cells, making the process even more nonstationary.

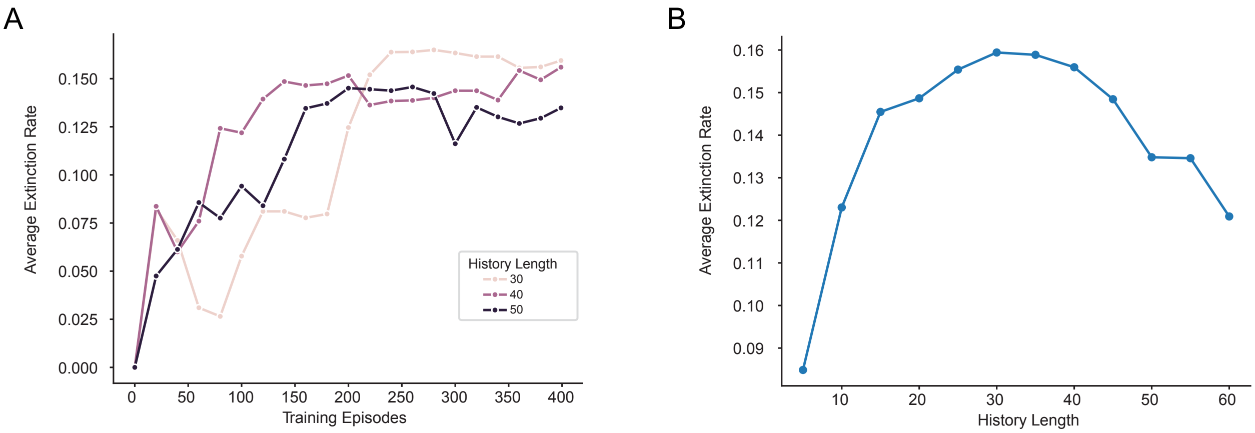


Fig. S1: **A** Performance dynamics, quantified by average extinction rate, across training episodes for three agents with different history lengths of past observations and actions. **B** Average extinction rate after 400 episodes of training for a range of agents, showing that there is a clear optimal history length (at 30 observations) which maximizes performance for a given training budget.

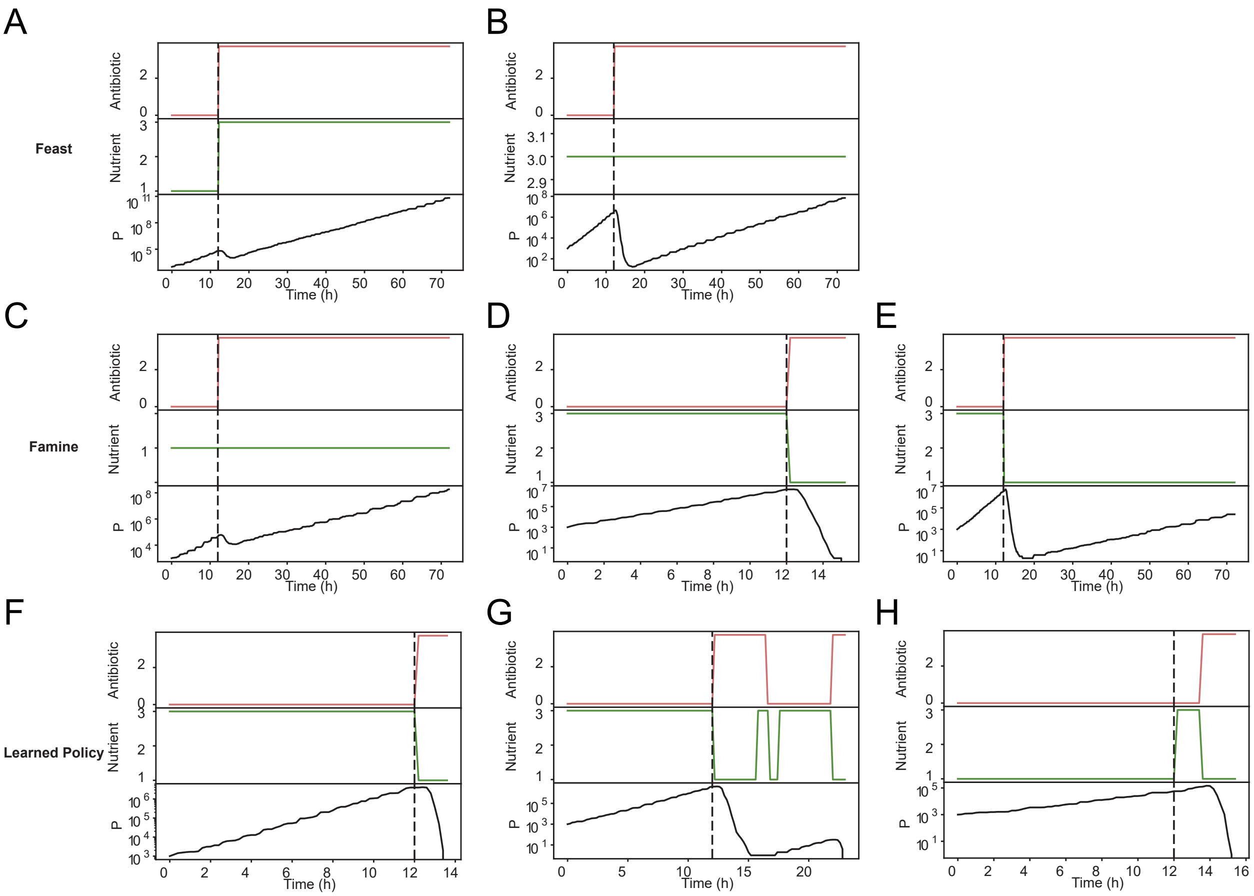


Fig. S2: **A-B** Additional example trajectories of the ‘Feast’ protocol, in which antibiotics are applied (top, red) concurrently with rich nutrients (middle, green), for different initial conditions (rich and poor nutrient quality). Regardless of initialization, this protocol fails to cause population extinction. **C-E** Additional example trajectories of the ‘Famine’ protocol, in which antibiotics are applied (top, red) concurrently with a nutrient-poor environment (middle, green), for different initial conditions (rich and poor nutrient quality). For populations initialized in rich nutrient environments, this approach results in heterogeneous population survival outcomes, with some realizations resulting in extinction, while others result in survival. **F-H** Additional example trajectories of the RL agent’s learned policy, in which antibiotics and nutrients are dynamically adjusted based on the population’s initial physiological state. Unlike the hard-coded policies, the agent is able to achieve extinction in every case by doing a famine-like protocol with treatment pauses in rich cells were re-exposed to rich nutrients.

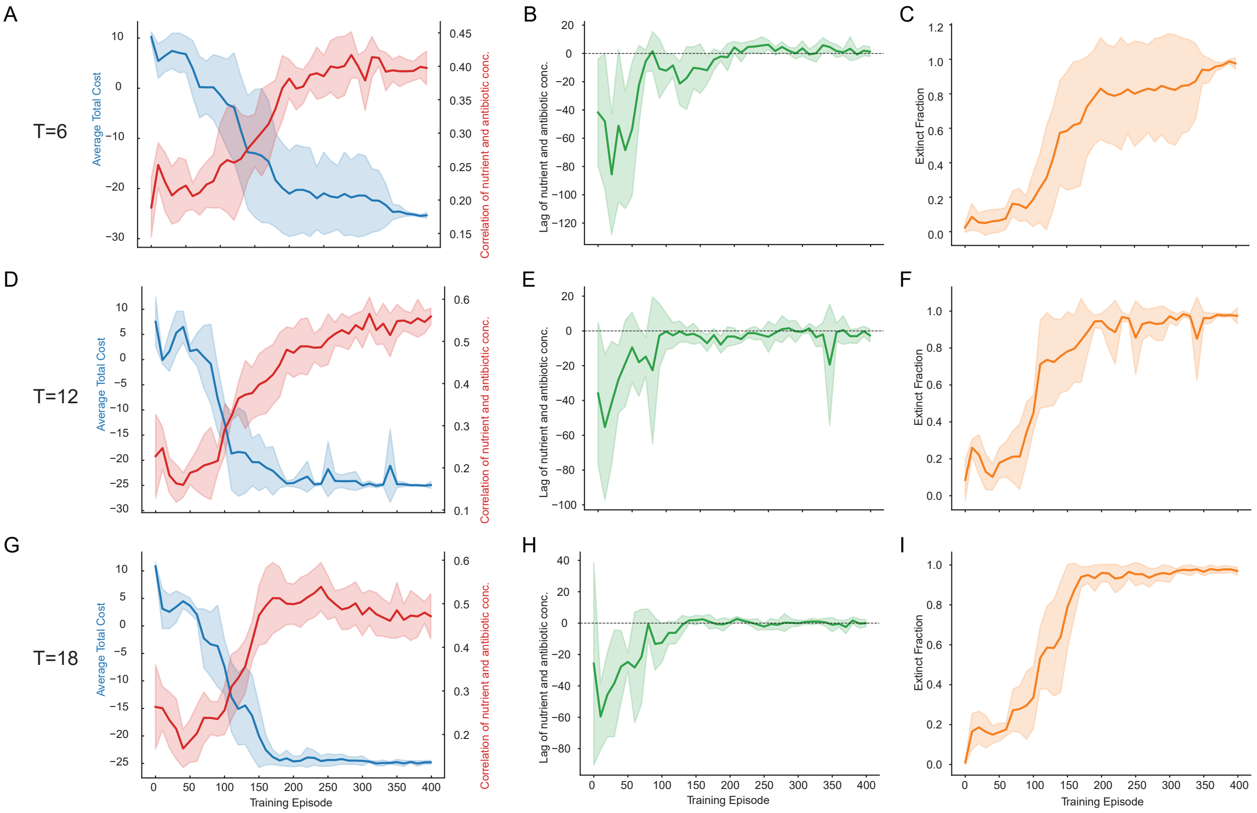


Fig. S3: **A,D,G** Average total cost (blue line), as quantified as the average log population ratio across an episode, and average cross correlation between nutrient and antibiotic timeseries (red line) across 400 training episodes, for different time periods of nutrient oscillation. Correlation between nutrient and antibiotic increases with performance, indicating the agent is able to infer the underlying nutrient state through training. **B,E,H** Lag of the maximum cross correlation between nutrient and antibiotic timeseries throughout training for different time periods, which stabilizes at a small positive value, indicating that the agent learns to respond to nutrient fluctuations when determining when to apply antibiotic regardless of time period. **C,F,I** Average extinction fraction throughout training for different time periods, indicating that extinction fraction stabilizes near one at the same time that lag and cross correlation stabilize, indicating that learned policy performance is dependent on successfully inferring the environmental state.

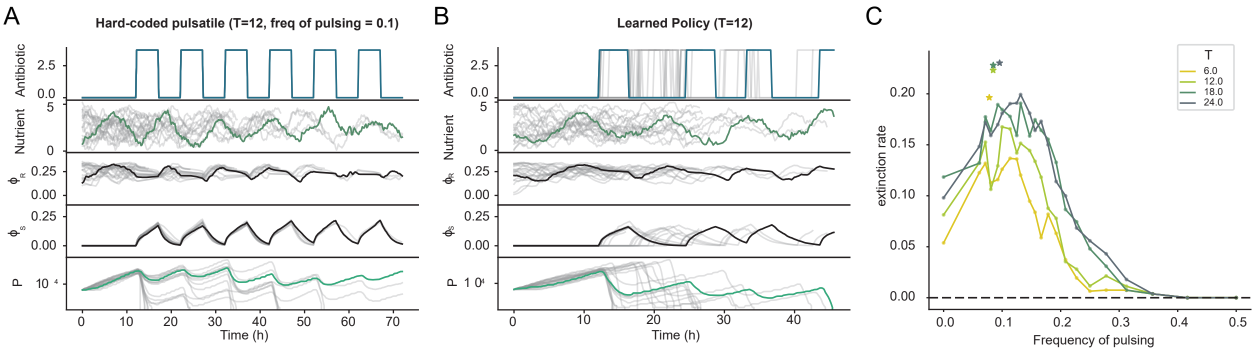


Fig. S4: Feedback-guided learned policies outperform hard-coded pulsing. **A,B** Agents were trained in fluctuating nutrient environments with $T = 12$ (**B**) and then evaluated and compared to a range of hard-coded pulsing protocols with different frequencies (**A**). **C** The learned policies were able to achieve a higher average extinction rate, defined as the inverse of the extinction time, than the hard-coded ones by adapting the pulsing frequency in response to differences in individual population growth trajectories.

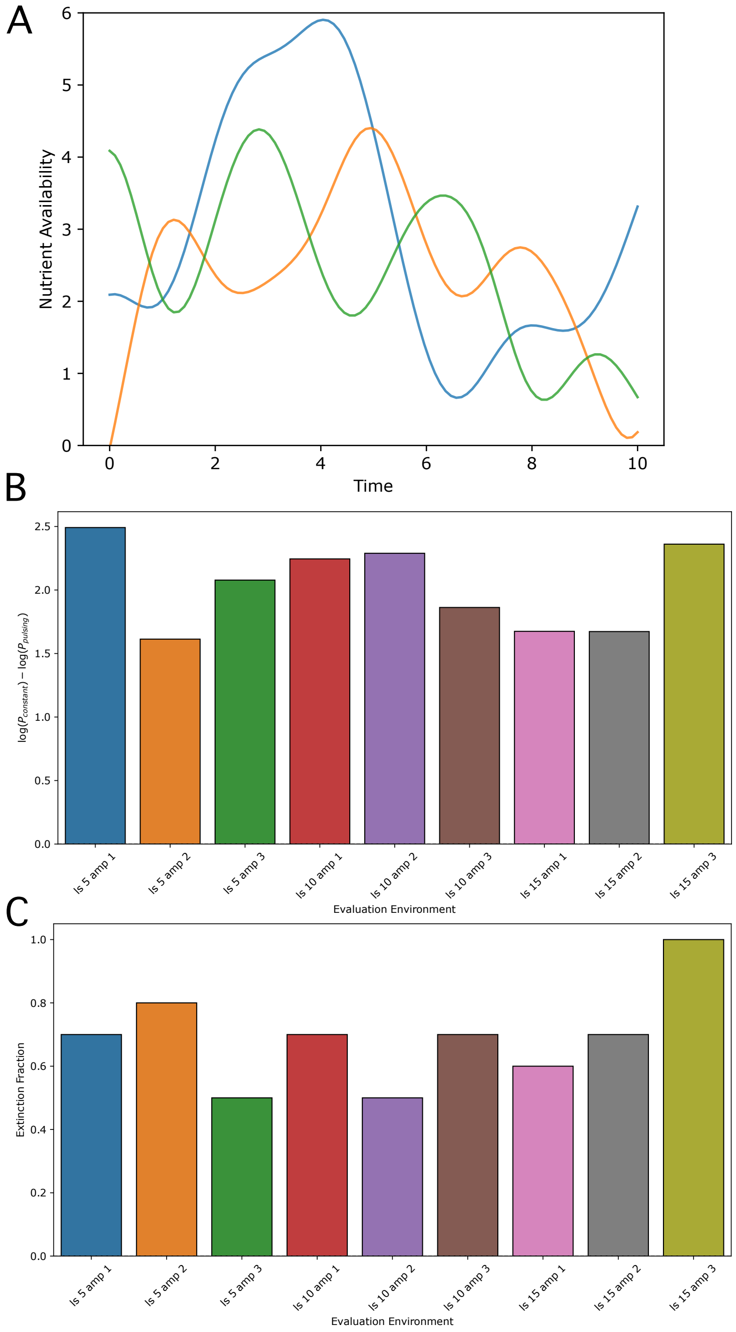


Fig. S5: Evaluation of the multi-environment trained agent on a variety of more complex fluctuating nutrient environments not seen in training. Specifically, the nutrient dynamics were defined by a Gaussian Process (GP) with RBF kernel: $k_{\mathrm{RBF}}(t, t') = (\mathrm{amp})^2 \exp(-||t - t'||^2/[2(\mathrm{ls})^2])$, where amp and ls denote the amplitude and length scale of the process, respectively. **A** Three representative GP simulations for amp = 1.5 and ls = 5. **B** Average log population reduction compared to constant drug application for 50 trials in different GP environments. **C** Average extinction fraction for the same environments. Agent performance generalizes well to all environments not seen in training, significantly outperforming constant application.
